# Retromer retrieves the Wilson Disease protein, ATP7B from lysosomes in a copper-dependent mode

**DOI:** 10.1101/2020.01.17.910125

**Authors:** Santanu Das, Saptarshi Maji, Ruturaj, Indira Bhattacharya, Tanusree Saha, Arnab Gupta

## Abstract

ATP7B utilizes lysosomal exocytosis to export copper from hepatocytes. We investigated the fate of ATP7B, post-copper export. At high copper ATP7B traffics to lysosomes and upon subsequent copper chelation, returns to Trans Golgi Network. At high copper, ATP7B co-localizes with lysosomal marker, Lamp1 and the core member of retromer complex, Vps35. Knocking down VPS35 did not alter copper-responsive vesicularization of ATP7B; rather upon subsequent copper chelation, ATP7B failed to relocalize to TGN that could be rescued by overexpressing wtVPS35. Using super-resolution microscopy and proximity ligation assays we demonstrate that VPS35 and ATP7B are juxtaposed on the same lysosomal compartment and their interaction is indirect. Utilizing in-cell photoamino acid-based UV-crosslinking and subsequent immunoprecipitation, we detected ATP7B and retromer subunits, VPS35 and VPS26 in a large complex in high copper conditions, hence confirming their interaction. We demonstrate that retromer regulates lysosome to TGN trafficking of the copper transporter ATP7B and it is dependent upon cellular copper level.

## Introduction

Lysosomes have traditionally been believed as a disposal organelle of the cell. A growing body of recent studies have implicated lysosomes as centers of cellular nutrient recycling (Abu-Remaileh et al., 2017; Korolchuk and Rubinsztein, 2011; Rabanal-Ruiz and Korolchuk, 2018). There are growing evidences that lysosomes regulate cellular homeostasis of various metals like, Cu, Zn and Fe. Polishchuk et al. (2014), Blaby-Haas and Merchant (2014), Kurz et al. (2011), Kambe (2011)). Copper, a transition metal serves as an essential micronutrient for biological system. It participates in redox reactions in different cellular metabolic pathways shuttling between Cu(II) and Cu(I) states (Uauy et al. (1998), where Cu^1+^ is favored for normal physiological activities (Fahrni (2013). Several proteins tightly regulate copper homeostasis and supplies bioavailable copper to the secretory pathway. Excess copper induces oxidative stress through Fenton reaction and hence is detrimental for living system (Gupta and Lutsenko (2009). Copper homeostasis is primarily maintained by two Trans Golgi Network (TGN) recycling P-type ATPase ATP7A (Menkes Disease Protein) and ATP7B (Wilson Disease Protein). ATP7A is expressed ubiquitously whereas expression of ATP7B is limited to liver, brain and kidney Telianidis et al. (2013) ATP7B solely functions to maintain the copper homeostasis in hepatocytes (Muchenditsi et al. (2017). Defects in ATP7B leads to Wilson Disease (WD), a phenomenon characterized with copper accumulation in liver, brain and other organs manifesting severe hepatic or neurological symptoms (Huster and Lutsenko (2007).

In this study we have attempted to dissect the trafficking itinerary of ATP7B and its mode of regulation in hepatocytes. At physiological/ basal copper level, ATP7B primarily resides on membrane of Trans Golgi Network (TGN) and functions in secretory pathway by delivering copper to Cu dependent ferroxidase enzyme, ceruloplasmin (Lutsenko (2016) within the lumen of TGN. At higher intracellular copper it vesicularizes (Anterograde trafficking), sequestering copper inside vesicles and exports copper in lysosomes (Polishchuk et al. (2014). What is the fate of copper in the lysosome is still to be determined. It is possible that the entire copper in the lysosome is excreted out of the cell by exocytosis. Alternatively, lysosomes may act as storehouse of bioavailable copper that is tapped as per the requirement of the cell.

Since some recent studies have shown that membrane cargoes recycle back from lysosomes (Seaman (2007); Suzuki and Emr (2018a); Canuel et al. (2008), we examined the fate of ATP7B at the lysosome. We specifically asked (a) Is ATP7B degraded at the lysosome during its copper export activity and (b) If not, what is the mechanism that retrieves ATP7B from lysosomal compartments?

Over 300 mutations in ATP7B are associated with WD and are frequently used to understand regulation and structure-function correlation of ATP7B (Aggarwal et al. (2013); (Gupta et al., 2005), Ala et al. (2015), Ala et al. (2005), Abdelghaffar et al. (2008), Braiterman et al. (2014), Caca et al. (2001). These reported disease mutations affect the functioning of ATP7B either by affecting its copper transporting activity or its trafficking or both (Braiterman et al. (2014), Gupta et al. (2011). Mentionable is the N41S at N-terminus, a naturally occurring WD mutation, which localizes at the TGN and basolateral membrane in all copper conditions (Braiterman et al. (2009). Another disease variant, Arg875 in the A-domain fails to escape ER but can be rescued with copper supplementation (Gupta et al. (2011).

Several proteins govern trafficking and stability of ATP7B (Materia et al. (2012); Jain et al. (2014); (Gupta et al., 2018) that interact directly or indirectly with defined and conserved motifs of the protein. These motifs influence the directionality of cargo transport between organelles like Golgi-endosome-plasma membrane. Both ATP7A and ATP7B harbors endocytic di/tri-leucine motif in their C-terminus which facilitates their transport across various compartments (Petris et al., 1998), (Braiterman et al., 2011). Francis et al have demonstrated that di-leucine motif ^1487^LL^1488^ of ATP7A to be important for its retrieval from cell membrane (Francis et al. (1999). Similarly, the ATP7B di-leucine mutant ^1454^LL>AA^1455^ caused redistribution of ATP7B from TGN to plasma membrane and dendritic vesicles and loss of somatodendritic polarity in rat hippocampal primary neurons (Jain et al. (2014).

Although the anterograde pathway of ATP7B has been moderately characterized (Gupta et al. (2016), Gupta et al. (2018), the regulation which mediates its retrograde transport from lysosomes has been elusive. Recent studies have shown that retromer regulates retrieval or rescue of cargoes from endosomal and lysosomal compartments (Burd and Cullen, 2014; Gershlick and Lucas, 2017; Lucas and Hierro, 2017; Tammineni et al., 2017). Retromer is a highly conserved endosomal sorting complex, composed of core components, VPS35, 26 and 29 and variable components, Sorting Nexins (SNX) and WASH complex, that are involved in the retrieval and retrograde transport of endocytosed transmembrane proteins (cargoes) to the trans-Golgi network (TGN) or cell surface (Seaman, 2018); Suzuki et al. (2019). The ATP7B homologue, ATP7A, has been shown to be regulated by SNX27 (Cullen and Korswagen, 2012) (Steinberg et al., 2013). In rat primary neurons SNX27, a variable member of the retromer complex, rescues neuroligin 2 from lysosomal degradation (Binda et al., 2019). Similarly, SNX17 protects integrin from degradation by sorting between lysosomal and recycling pathways, though retromer is not directly involved. Recycling of CI-M6PR, the protein which delivers acid hydrolases to lysosome, to TGN is dependent on Retromer (Seaman, 2007). Studies from Emr group have shown that in yeast proteins like autophagy protein Atg27 is recycled from vacuole to the endosome via the Snx4 complex and then from the endosome to the Golgi via the retromer complex (Suzuki and Emr (2018b). Further they have also demonstrated that both VPS26 and VPS35 are critical in cargo retrieval; however VPS26 utilizes different binding sites depending on the cargo, allowing flexibility in its cargo selection (Suzuki et al. (2019). Besides TGN delivery, retromers also regulate endosome-to-plasma membrane recycling as in Ankyrin-repeat domain 50, ANKRD50 (McGough et al. (2014).

In this study, we have demonstrated that the recycling Cu-P-type ATPase, ATP7B, whose localization in the cell is dictated by intracellular copper levels, recycle from lysosome to TGN upon copper removal and this phenomenon is regulated by the retromer. We have demonstrated that similar to as in case of an endocytic cargo (Mellado et al. (2014), Tabuchi et al. (2009) retromer also sorts a secretory cargo, i.e., ATP7B for its TGN delivery from lysosomes and late endosomes.

## Results

### ATP7B recycles between Lysosomes and Trans Golgi Network in a copper dependent manner

ATP7B vesicularizes from trans-Golgi network in response to high copper. To determine the optimal time of complete retrieval of ATP7B from vesicles to TGN, HepG2 cells (human hepatocellular carcinoma cell line) were treated with high copper (50μM; 2h) and subsequently treated with 50μM BCS for varying time periods (10min, 30min and 2h). Gradual increase in colocalization between TGN and ATP7B was observed with 10 min, 30 mins and 2h of BCS treatment as evident from Pearson’s Colocalization Coefficient (PCC). At 2h, maximum TGN retrieval of ATP7B was observed (Fig. 1, A and B).

**Fig1:**
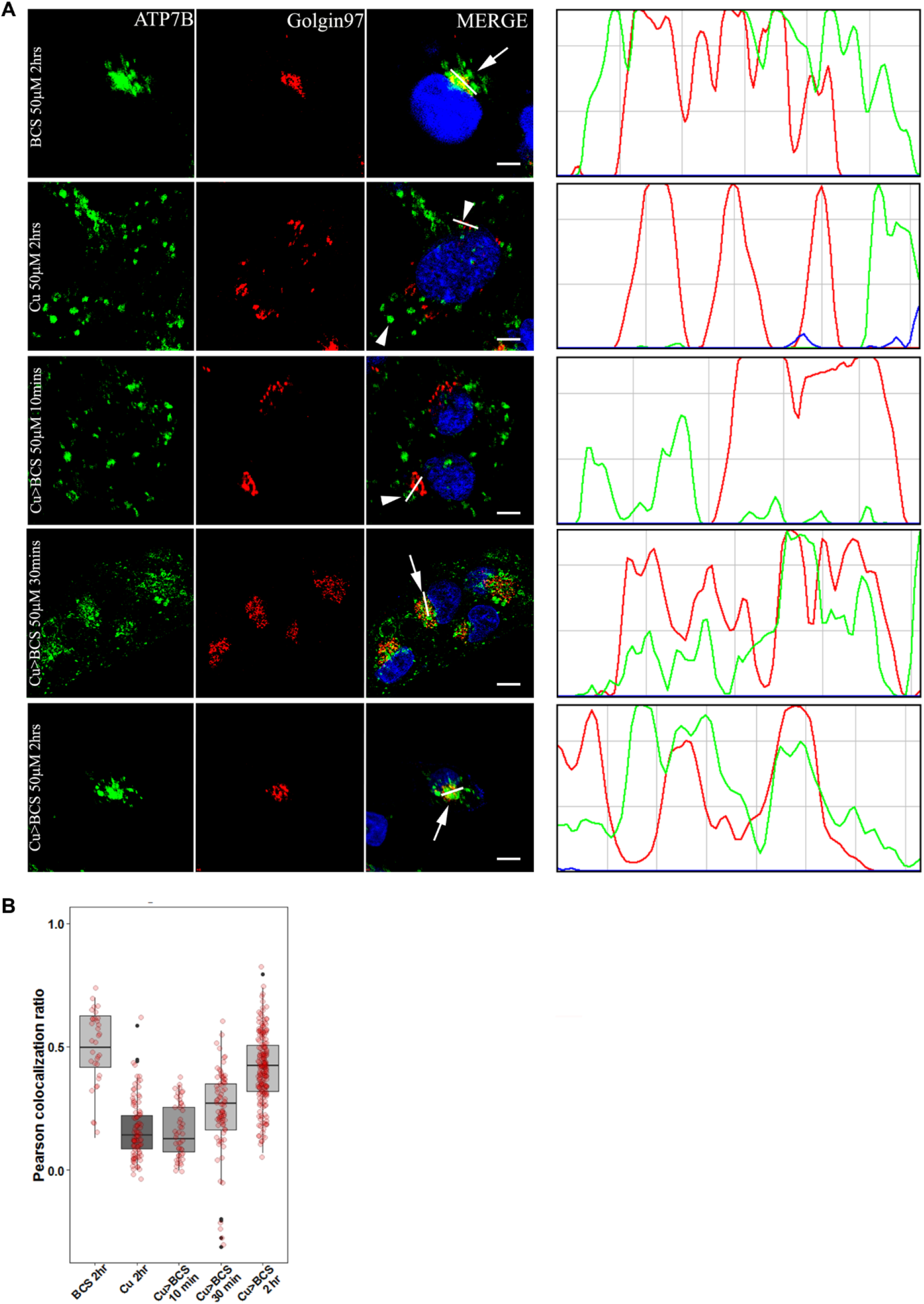
ATP7B recycles between TGN and vesicles in a copper dependent manner: (A) Colocalization of ATP7B (green) with TGN marker, Golgin97 (red) in copper limiting, BCS (top panel) and 50μM copper (panel 2). Panel 3-5 shows subsequent return of vesicularized ATP7B upon BCS treatment for varying time length (10, 30 and 120mins). The overlap plots (right boxes) show the extent of overlap of green and red at lines drawn through the signals located on TGN (marked by arrow). Arrow and Arrowhead represents TGN localized and vesicularized ATP7B respectively. Scale bars represents 5μM. Blue signal represents DAPI staining for nucleus. (B) Pearson’s correlation coefficient of colocalization between ATP7B and Golgin97 at different copper conditions demonstrated by a box plot with jitter points

Polishchuk et al have demonstrated that ATP7B utilizes lysosomal exocytosis to export copper (Polishchuk et al., 2014). We investigated whether ATP7B degrades at the lysosome after it transports copper or it recycles back from lamp1 positive lysosomal compartments for a next round of export cycle. We treated the cells with either BCS (50μM) or increasing amounts of copper (50μM and 250μM). Using immunofluorescence assay we determined that under high copper conditions ATP7B exits TGN and colocalizes with Lamp1 and to a lesser extent with Rab7 positive compartments (Fig. 2, A and B and Fig. S1, A and B). Upon treatment with 250uM copper, a drop of ATP7B protein abundance was observed compared to the two other experimental conditions indicating possible degradation (Fig. 2, C and D). Upon triggering the retrograde pathway with BCS for 30 mins (50μM), for cells that were pre-treated with 50uM copper, we noticed loss of colocalization of lamp1 and ATP7B, indicating lysosomal exit of ATP7B upon copper chelation (Fig. 2A). We did not observe appreciable colocalization of recycling marker, Rab11 and ATP7B (Fig. S1C). Interestingly irrespective of copper treatment we observed a fraction of ATP7B consistently colocalizing with lamp1.

**Fig2:**
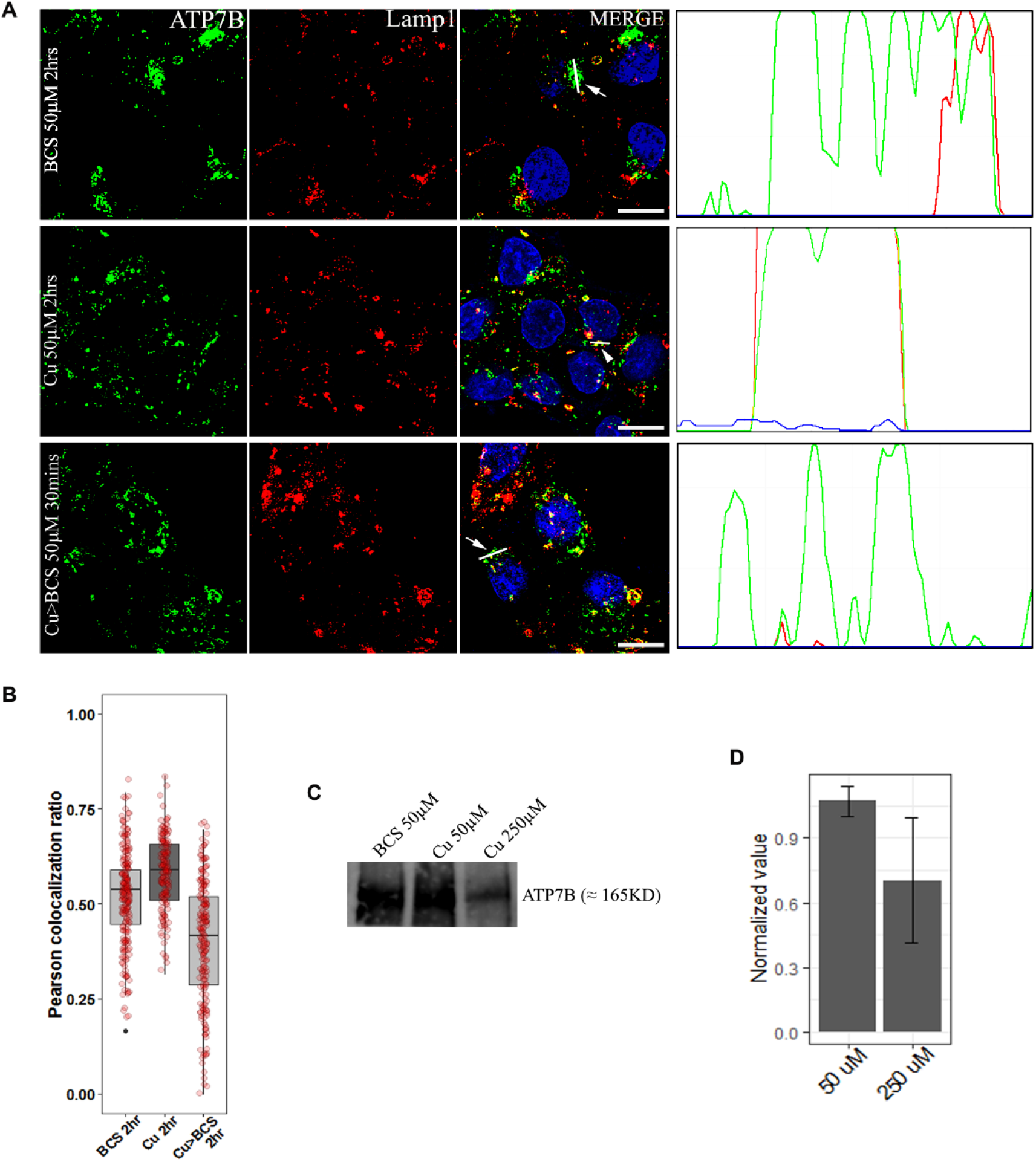
ATP7B recycles from lysosome upon copper depletion: (A) Colocalization of ATP7B (green) with lysosomal marker, lamp1 (red) in copper limiting, BCS (top panel) and 50μM copper (panel 2) and copper depletion post copper treatment (bottom panel). The overlap plots (right boxes) show the extent of overlap of green and red at lines drawn through the signals (marked by arrow or arrowhead). Arrowhead represents vesicularized ATP7B and arrow represents perinuclear positioned ATP7B. Scale bars represents 5μM. Blue signal represents DAPI staining for nucleus. (B) Pearson’s correlation coefficient of colocalization between ATP7B and Lamp1 at different copper conditions demonstrated by a box plot with jitter points. (C) and (D) Comparative protein abundance of ATP7B (normalized against total membrane protein) determined by immunoblot at different copper and copper-depleted conditions (Cu: 50 μM and 250 μM and BCS: 50 μM).

Lamp1 typically marks the terminal end of the endo-lysosomal pathway (Humphries et al. (2011), but frequently, Rab7 and Lamp1 also co-labels the common non-degradative lysosomal compartment (Cheng et al., 2018). To determine the percentage distribution of ATP7B in the milieu of lysosomes-late endosomal compartments (lamp1-Rab7), we utilized Structured Illumination Microscopy (SIM) along with deconvolution confocal microscopy imaging. At elevated copper (50μM) we found a proportionately higher co-distribution of ATP7B with compartments positive for both Rab7 and Lamp1 than compartments positive for individual unique markers. At this condition, 48.5% ± 13.2 of total ATP7B colocalized with either Lamp1 or Rab7 or both. Among them, 44.9% ± 14 of ATP7B localized in vesicles positive for both Lamp1 and Rab7. Hence, it may be inferred that recycling of ATP7B to TGN upon Cu chelation is largely from non-degradative transitory type vesicles which are positive for both the markers than classical lamp1-only lysosomal compartments (Fig. 3, A and B). Together this result suggests that ATP7B localizes at late endo-lysosomal compartments at high copper. However, at this point we could not determine whether the retrograde pathway of ATP7B can also originate exclusively from lysosomal or late endosomal compartments.

**Fig3:**
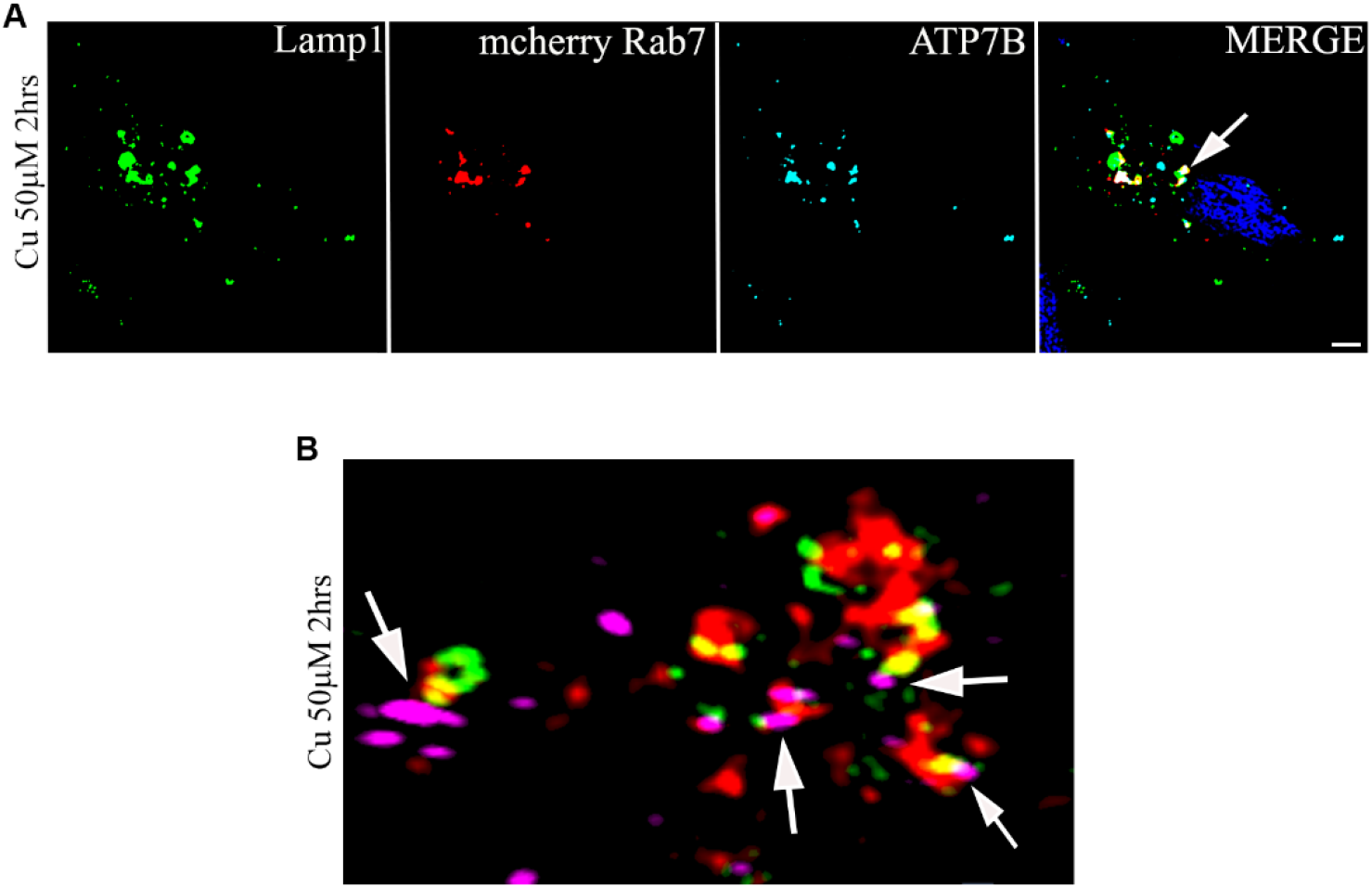
ATP7B is preferentially located in Rab7-Lamp1 positive endosomes in high copper: (A) Colocalization of ATP7B (cyan) with lysosomal marker, lamp1 (green) and late endosome marker, mCherry Rab7 (red) at 50μM copper. (B) 3D representation of Structured Illumination Microscopy (SIM) image of same with 100nm resolution. ATP7B is marked in magenta, Lamp1 in green and mCherry Rab7 in red. Arrow represents triple merging. Blue signal represents DAPI staining for nucleus.

It is worth mentioning that the size and shapes of the lysosomal compartments (Lamp1 +) are variable across cells and as well as in a single cell. It varies from 20-200μM in diameter (maximum length across) and shape varies from small puncta to larger patchy clumps. We did not notice any correlationship between size and shape of the compartment with the treated copper concentration or the passage number of the cell.

### Retromer regulate retrograde trafficking of ATP7B from lysosome and late endosome

ATP7B exits lysosomes by an unknown regulatory mechanism upon copper chelation. It has been shown that Menkes disease protein, ATP7A, the homologue of ATP7B require SNX27-retromer to prevent lysosomal degradation and maintain surface levels and localization (Steinberg et al. (2013). Also, CI-M6PR (a constitutive lysosomal recycling cargo) recycles to TGN from lysosomes in a retromer regulated fashion (Arighi et al., 2004) (Cui et al., 2019). This prompted us to investigate if the retromer complex plays a role in retrieval of ATP7B (a non *bona fide* lysosomal cargo) from lysosome and late endosomal compartments. VPS35 is the largest core component of the retromer complex. It functions as the scaffold for the assembly of other core components, VPS29 and VPS26 and also cargo binding (Hierro et al. (2007). Hence, we selected hVPS35 as the target component to determine the role of retromer complex, if any in copper mediated trafficking of ATP7B.

Broadly, localization and trafficking of ATP7B in HepG2 can be divided in 4 *phases* (a) at the TGN in Basal Cu (or–Cu), (b) on the anterograde vesicles in high Cu and (c) on the retrograde vesicles in high Cu > Cu chelated conditions (10 mins BCS treatment) and (d) majority of ATP7B back to the TGN in high Cu > Cu chelated conditions (30 mins/2h BCS). Immunoblot analysis of VPS26 and VPS35 revealed that HepG2 cells expresses the retromer complex proteins (Fig. 4A) and copper does not alter abundance of VPS35 in HepG2 cells (Fig. S2A)

**Fig4:**
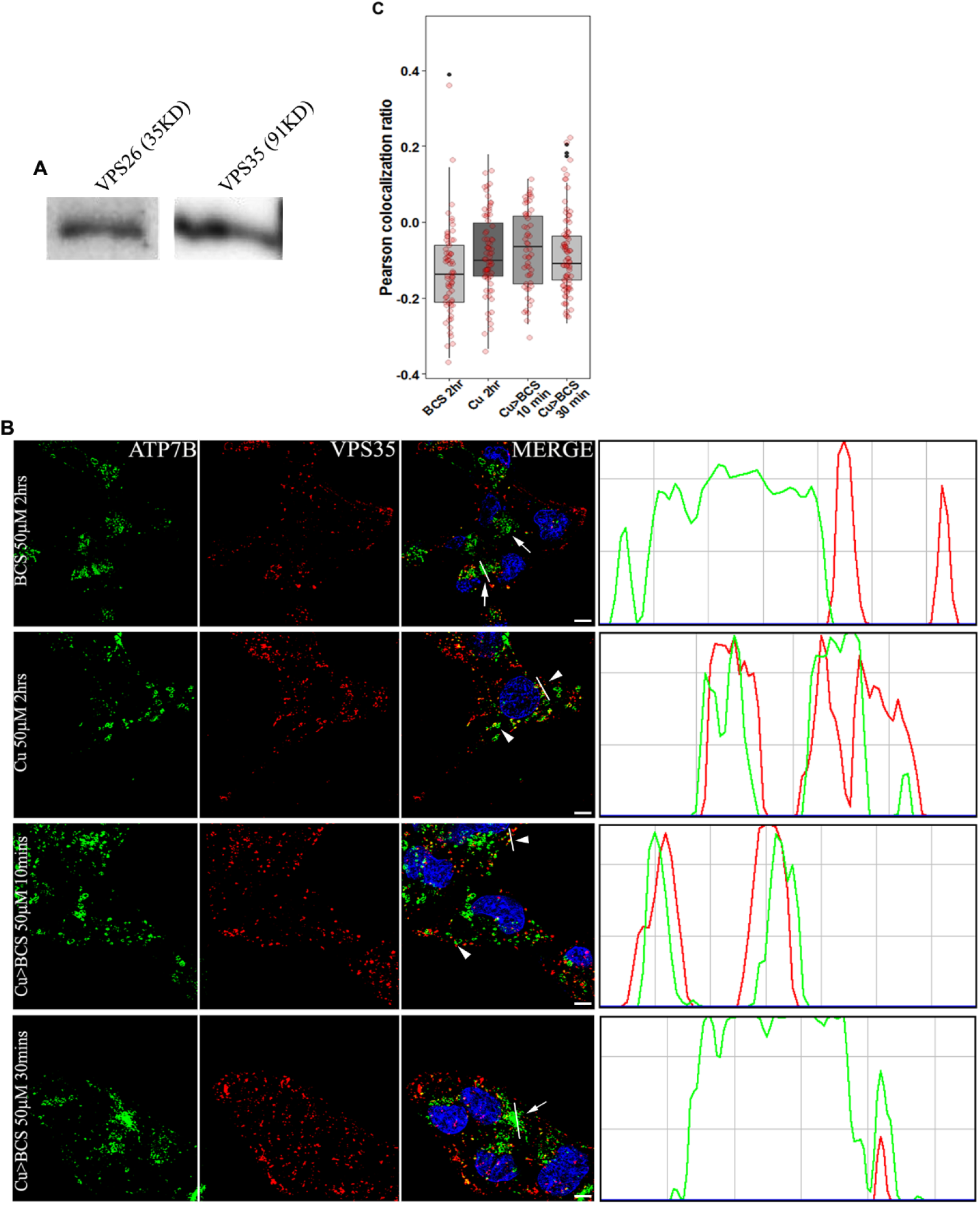
ATP7B and VPS35 colocalizes at high copper: (A) Immunoblot showing HepG2 expresses the retromer subunits VPS26 and VPS35 (B) Colocalization of ATP7B (green) with retromer subunit, VPS35 (red) in copper limiting, BCS (top panel) and 50μM copper (panel 2) and copper depletion post copper treatment (panels 3-4). The overlap plots (right boxes) show the extent of overlap of green and red at lines drawn through the signals (marked by arrow or arrowhead). Arrowhead represents vesicularized ATP7B and arrow represents perinuclear ATP7B. Scale bars represent 5μM. Blue signal represents DAPI staining for nucleus. (C) Pearson’s correlation coefficient of colocalization between ATP7B and VPS35 at different copper conditions demonstrated by a box plot with jitter points

To study if ATP7B co-localizes with retromer core components, cells were treated with either of the 4 conditions described above. Cells were fixed, blocked and co-stained with anti-ATP7B and anti-VPS35 or anti-VPS26 antibody. Maximum colocalization as quantified by Pearson’s Colocalization Coefficient between VPS35 and ATP7B was observed in Cu (2h)>BCS (10min) (*phase c*) followed by high copper condition (*phase b*) condition. Further with 30 min BCS treatment post high copper (*phase d*), ATP7B and VPS35 shows loss of colocalization (Fig. 4, B and C). Similar observations were made with VPS26 (data not shown).

To further understand if retromer regulates any of these phases of ATP7B trafficking, we knocked down VPS35 and studied the phenotype of copper induced localization of ATP7B with respect to TGN. Appreciable knockdown was attained (>70-80%) for the targeted siRNAs as compared to scrambled for VPS35 as ascertained by immunoblotting (Fig. 5A). Furthermore, as reported by (Fuse et al. (2015) decrease in expression of VPS26 was also observed in VPS35 KD cell which eventually tells us expression as well as functionality of VPS subunits are interdependent (Fig. S2B).

**Fig5:**
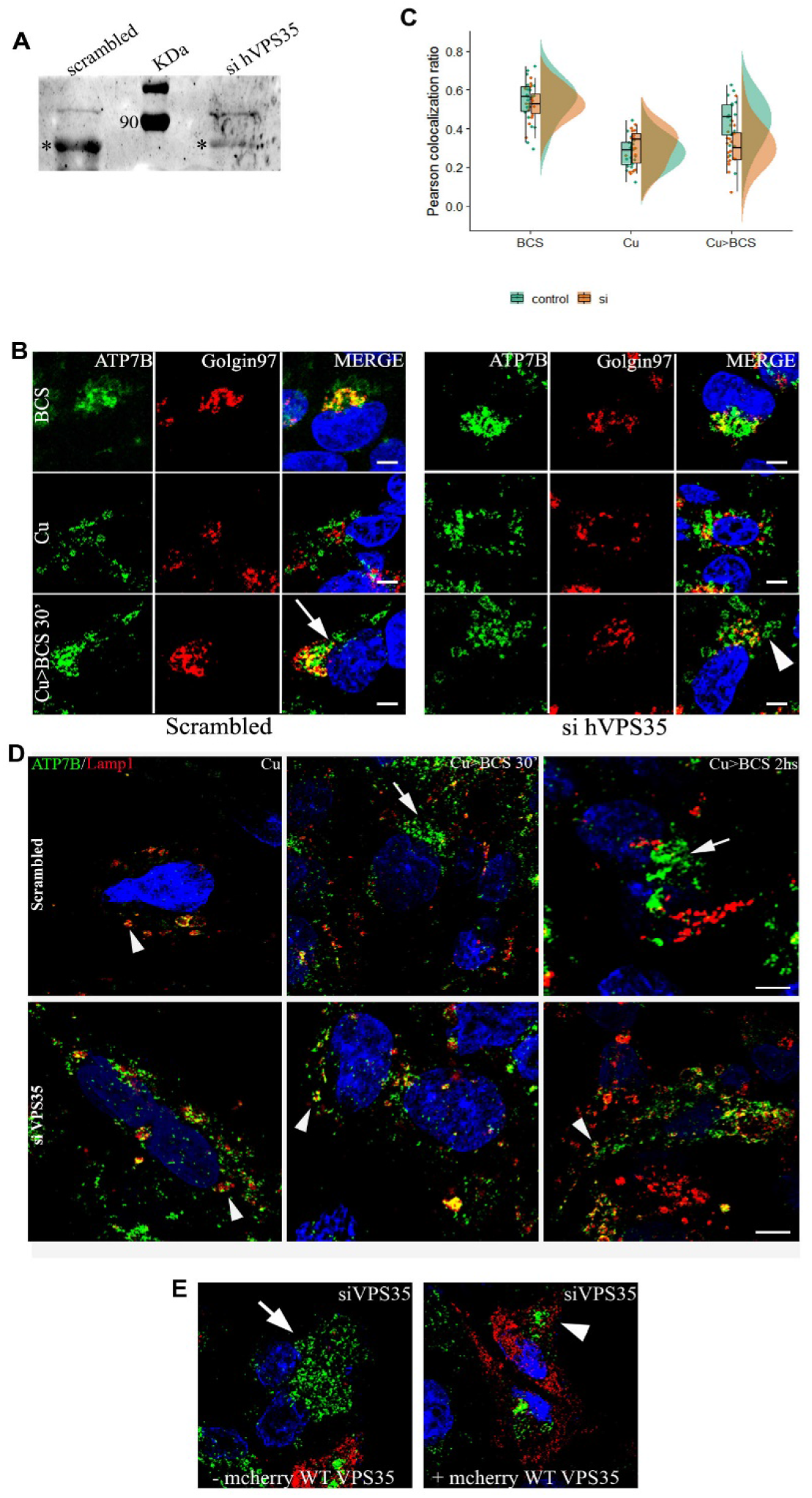
VPS35 regulates retrieval of ATP7B from lysosomes to TGN: (A) siRNA mediated knockdown of Vps35 in HepG2 cells shows its downregulation. (*) denotes the VPS35 protein (B) Colocalization of ATP7B (green) with TGN marker, Golgin97 (red) in BCS (top panel) and 50μM copper (panel 2) and copper depletion post copper treatment (panels 3). Arrow denotes TGN colocalization of ATP7B, Arrowhead denotes vesicularized ATP7B. Scale bars represent 5μM. (C) Pearson’s correlation coefficient of colocalization between ATP7B and TGN at different copper conditions comparing VPS35 siRNA treated vs control demonstrated by a box plot. (D) Merged image showing colocalization of ATP7B (green) with Lamp1 (red) in high copper (top/bottom left) and copper depletion post copper treatment for 30 mins and 2h (top/bottom middle and right respectively). The top panel represents control cells transfected with scrambled siRNA and bottom panel represents cells with VPS35 siRNA. Arrows mark perinuclear ATP7B and arrowheads denote vesicular ATP7B (E) Localization of ATP7B (green) in VPS35 siRNA treated cells and subsequently transfected with mCherry-wtVPS35 (red). The left image represents cells that is not expressing mCherry-wtVPS35 (arrow represents vesicularized ATP7B) as compared to cells expressing the construct (arrowhead represents presence of tight perinuclear ATP7B). Cells belong to the same culture dish for both the images. Blue signal represents DAPI staining for nucleus.

VPS35 siRNA treated cells were incubated with (a) BCS or (b) Copper or (c) Copper > BCS (30 min), fixed, blocked and stained with the anti-ATP7B and anti-Golgin97 antibodies. We observed that trafficking of ATP7B from the vesicles back to the TGN (*condition c*) was significantly abrogated (Fig. 5, B and C and Fig. S2C). ATP7B remained in vesicles and failed to recycle back to TGN even after the cells were incubated in BCS for a prolonged time of 2h subsequent to copper treatment (not shown).

To corroborate our finding that VPS35 regulates ATP7B trafficking, we utilized the wild type and the inactive dominant negative mutant mCherry-VPS35 (R107A) (gift from Dr, Sunando Datta, IISER Bhopal). The R107A mutation abolishes the interaction of VPS35 with VPS26 and affects cargo sorting. (Gokool et al. (2007); Zhao et al. (2007). Cells were co-transfected with GFP-ATP7B and mCherry-wt-VPS35 and GFP-ATP7B was concentrated at TGN (perinuclear region) with copper chelator. Vesicularization and recycling of ATP7B was triggered with treatment with copper. Image capture was initiated at the point of copper treatment and data was collected at an interval of 1.93 second for a total period of 30 mins. We noticed that within 1 min of Cu treatment, GPF and m-Cherry signals colocalized at the same endosomal vesicle. The dwell time of wt-VPS35 and GFP-ATP7B on the vesicle was significantly higher for wt-VPS35 compared to the mutant. For VPS35-R107A, the colocalization lasted for few seconds (4-11 seconds) but for wt-VPS35, the colocalization lasted for an average of 7 mins. (Video 1, A and B) and (Fig. S3, A and B).

Since we determined that ATP7B colocalizes at the lysosomes at high copper, we investigated if VPS35 regulates lysosomal exit of ATP7B on triggering its retrograde pathway. Using identical experimental conditions of knocking down VPS35, we found that ATP7B is arrested at lysosome upon triggering the retrograde pathway (i.e., high copper > BCS) (Fig. 5D). Interestingly, boosting the retrograde pathway by lengthening BCS treatment time to 2h did not facilitate retrieval of ATP7B from the lysosomes (Fig. 5D). Also, in VPS35 kd cells, a population of ATP7B was arrested in late endosome (Rab7 positive) upon activating the retrograde pathway (Fig. S2C).

We confirmed the role of retromer by rescuing the non-recycling phenotype of ATP7B in VPS35 kd cells by overexpressing mcherry-wt-VPS35. We found that ATP7B recycled back from vesicular to its tight perinuclear TGN localization upon copper chelation in VPS35 kd cells that overexpressed the wt-VPS35 construct (Fig. 5E). These experiments confirm that VPS35 regulates retrieval of ATP7B from lysosomes (and also possibly from late endosomes) to TGN upon copper depletion.

### Lysosomal luminal pH does not influence localization of ATP7B and recruitment of VPS35

It emerges that copper induced localization of ATP7B involves a tripartite participation, i.e., ATP7B, lysosome and retromer. After confirming the role of VPS35 in this process, we asked if luminal lysosomal environment affect retromer recruitment and hence ATP7B retrieval from lysosome. Retromers have been previously implicated in lysosomal activity e.g., autophagy (Cui et al. (2019). We investigated if the targeting of ATP7B to lysosomes in high copper or its retrieval initiation by VPS35 recruitment is affected by inactivation of the V-ATPase that is crucial for lysosomal functioning. Cells were treated with the V-ATPase inhibitor, BafA1, in 50μM copper (to trigger lysosomal targeting of ATP7B) and Cu>BCS (20 mins) conditions (to trigger its lysosomal exit). No observable and significant difference in colocalization of Lamp1, ATP7B and VPS35 was obtained between BafA1 treated vs the control in either condition. It can be inferred that retromer being situated on the outer membrane of the Lamp1 positive compartments is unaffected by the change in luminal pH of the lysosome brought on by BafA1 treatment. Localization of ATP7B also stays unaltered as also demonstrated by Polishchuk et al. (Fig. 6)

**Fig6:**
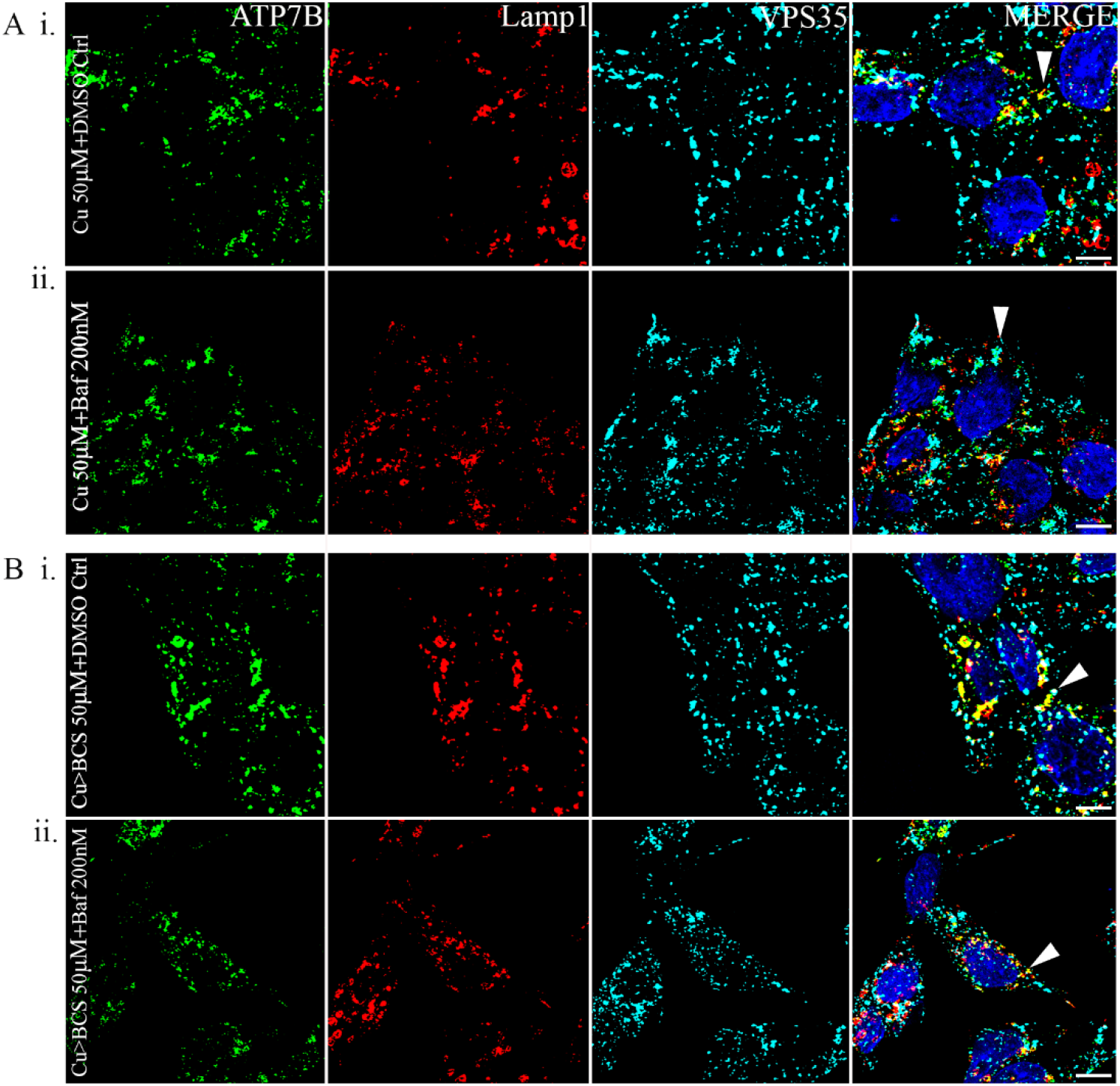
Lysosomal luminal pH does not influence localization of ATP7B and recruitment of VPS35: (A) Colocalization of ATP7B (green), Lamp1 (red) and VPS35 (cyan) in high copper for 2hrs in cells treated with Bafilomycin 1 (lower panel) or not (upper panel). (B) Colocalization of ATP7B (green), Lamp1 (red) and VPS35 (cyan) in cells treated with copper chelator for a brief period (20 mins) subsequent to high copper treatment to induce ATP7B vesicularization. Cells were treated with Bafilomycin 1 (lower panel) or not (upper panel).

### VPS35 acts on ATP7B in a micro-distant *modus operandi*

Next, using biochemical assays to investigate if VPS35 directly interacts with ATP7B we utilized co-immunoprecipitation where GFP-ATP7B was expressed in cells, pulled down with anti-GFP beads and probed for endogenous VPS35. We failed to detect any signal in the immunoblot developed with anti-VPS35 antibody (Fig. S4).

To understand the underlying reason of why we could not detect interaction between ATP7B and VPS35 using biochemical methods, we resorted to Super resolution microscopy to determine the exact positioning of ATP7B w.r.t VPS35 at the lysosomal compartment. Using Structured Illumination Microscopy and High resolution deconvolution confocal microscopy we observed that ATP7B and VPS35 lies in juxtaposition on the lysosomal compartment (stained with Lamp1 antibody) at high copper conditions (Fig. 7, A-D). The average distance between these two proteins varies from 25nm~200nm. It can be inferred that although ATP7B lies in close proximity to VPS35 and is regulated in its retrograde pathway by retromer, the physical interaction between these two proteins are indirect. We utilized Stimulated Emission Depletion (STED) microscopy to look further closely on the disposition of ATP7B and Vps35 on a vesicular membrane. Using Z-stacking we determined the shape of a vesicle (dotted circle in Fig. 7E) and observed that ATP7B (green) and Vps35 (red) decorated the vesicular membrane with minimal signal overlap (yellow) at a maximum resolution of 25nm further substantiating our biochemical findings.

**Fig7:**
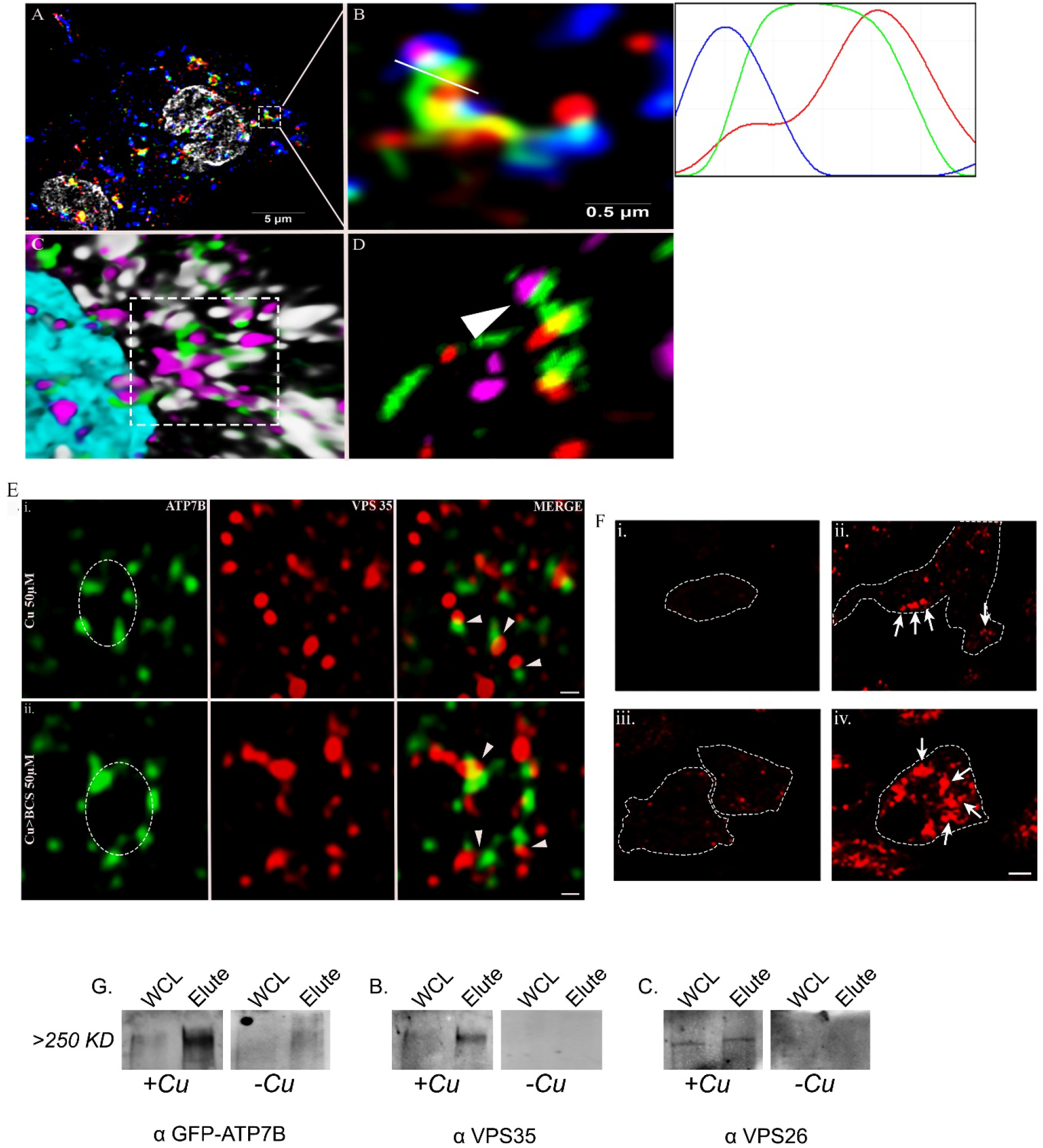
VPS35 interacts with ATP7B on lysosome in a micro-distant manner: (A) High resolution deconvoluted confocal microscopy merged image showing colocalization of ATP7B (green) with Lamp1 (red) and VPS35 (blue) at 50μM copper. Grey represents nucleus. (B) Zoomed image of inset in A. The overlap plots (right box) show the extent of overlap of green, red and blue at lines drawn through the signals (marked by white line). (C) 3D representation of zoomed image in B, marked by dashed line. ATP7B is marked in green, Lamp1 in grey, VPS35 in magenta. Cyan represents nucleus. (D) 3D representation of Structured Illumination Microscopy (SIM) image of same with 100nm resolution. ATP7B is marked in green, Lamp1 in red and VPS35 in magenta. Arrowhead represents co-distribution of ATP7B and VPS35 in lysosomal compartment (Lamp1). (E) Stimulated emission depletion (STED) microscopy image of ATP7B (green) and VPS35 (red). Top panel and bottom panel shows colocalization of ATP7B and VPS35 in high copper and copper depletion post copper treatment respectively. In both conditions ATP7B containing vesicles (marked by dotted circle) show juxta-positioning of VPS35 (red) and ATP7B (green). Arrowhead represents point of juxtaposition or merging. Scale bars represents 200nM. (F) Proximity Ligation Assay: *Upper panel* shows interaction of mouse Hur and rabbit Trim21 in HepG2 which serves as a positive control; **i.** Technical negative control (NC) without primary antibodies probed with only anti-rabbit(−) and anti-mouse(+) secondary PLA antibodies. **ii.** Hur and Trim21 interaction, marked by arrows. *Lower panel* shows interaction of rabbit ATP7B and goat VPS35 in HepG2; **iii.** Technical negative control (NC) without primary antibodies probed with only anti-rabbit(−) and anti-goat(+) secondary PLA antibodies. **iv.** ATP7B and VPS35 interaction, marked by arrows. (G) In-cell crosslinking A. Immunoblot showing presence of GFP both in WCL and elute in +Cu state at ~ >>250KDa (metabolically crosslinked with photo-amino acids) which was absent in −Cu state. B. and C. Similar pattern was observed for both co-eluted proteins VPS35 and VPS26 respectively; *WCL: Whole cell lysate*

Since, the interaction between ATP7B and VPS35 is likely to be indirect and possibly they a part of a larger complex, we utilized **P** roximity **L** igation **A** ssay (PLA) to substantiate our finding. Interaction of Hur and Trim21 (Guha et al., 2019) were used as used as positive controls and secondary antibodies against ATP7B (anti-rabbit) and VPS35 (anti-goat) were used as negative controls. We observed positive interaction between ATP7B and VPS35 that is evident by formation of red intracellular puncta on sites if the two proteins juxtapose at a distance less than <40nm (Fig 7F).

This finding strongly suggest that VPS35 and ATP7B are a part of a larger complex and the regulation of ATP7B trafficking by the retromer proteins are possibly indirect and they do not share a physical interface. To detect such an interaction, we metabolically labeled GFP-ATP7B overexpressing HepG2 cells with Photo amino-acids (P-Leucine and P-Methionine). Cells were treated with copper or copper chelator and subjected to UV crosslinking. Crosslinked GFP-ATP7B complex was immune-precipitated with anti-GFP beads. Upon probing the eluate on an immunoblot with anti-VPS35, VPS26 and anti-GFP antibodies, a complex >> 250KDa was detected that was comprised of all the three proteins. The complex was found to be absent or much less abundant in cells treated with copper chelator, TTM (Fig 7G).

In summary we establish that, ATP7B traffics to lysosomes at high copper where it juxtaposes with VPS35 and contributes to the formation of a complex that comprises of at least ATP7B, VPS35 and VPS26 in addition to other proteins. Upon triggering the retrograde pathway by subsequent copper chelation, retromer regulates the recycling of ATP7B from the lysosome to the TGN (Fig. 8).

**Fig8:**
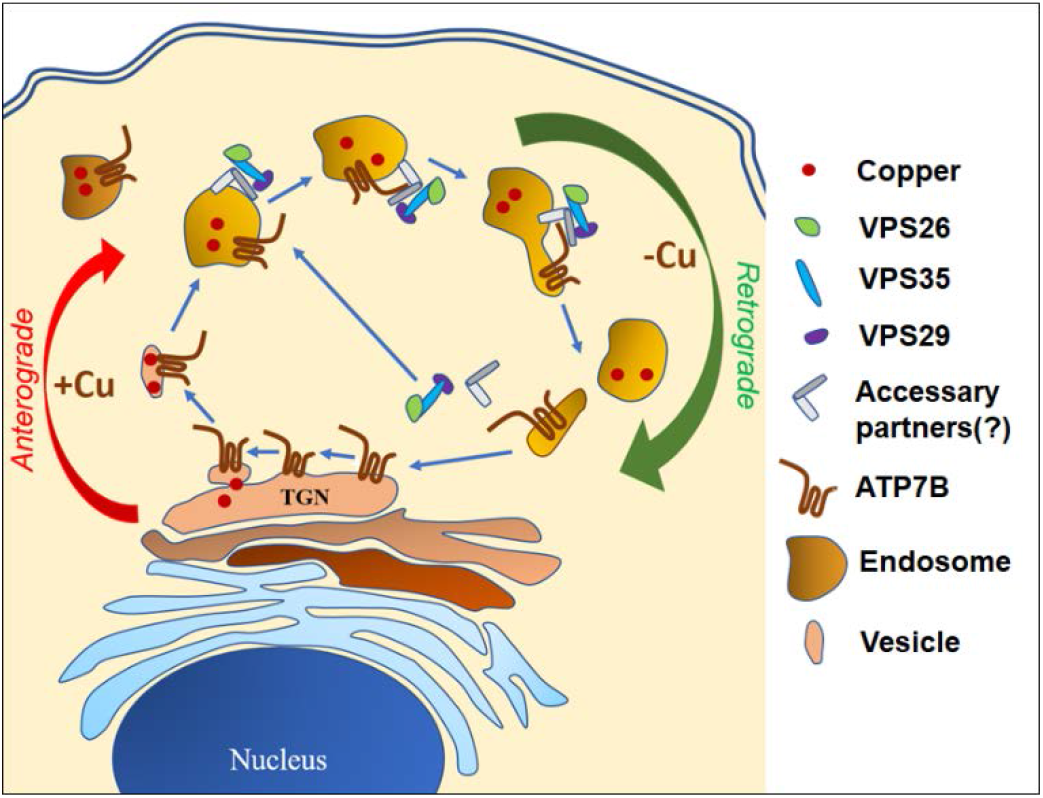
Schematic representation of recycling of ATP7B between TGN and lysosomes. Retromers are recruited on the lysosomal membrane that regulates retrograde transport of ATP7B upon copper removal.

## Discussion

The copper transporting ATPase, ATP7B exports copper through lysosomes. ATP7B (160kDa) is a large 8 membrane spanning protein with a total of 1465 residues. ATP7B resides on the TGN membrane at basal copper levels and traffics to lysosomes and Rab7 compartments at high copper. We argue that it would be highly wasteful for the cell to degrade ATP7B after each cycle of copper export from the TGN to the lysosomes. We wondered whether ATP7B recycles back from the lysosome after it pumps copper in the lysosomal lumen for either export out of the cell or for reutilization as a nutrient that requires to be investigated. Interestingly, the protein does not get degraded unlike most other cargoes that are destined for degradation at the lysosomes. The luminal loops of ATP7B between the 8 TM domains are small and probably escapes lysosomal hydrolases. Also, low pH in lysosomal lumen might help the release of copper from the His residues that are located between TM1-TM2 loop as shown in its homologue ATP7A (Barry et al., 2011; LeShane et al., 2010; Otoikhian et al., 2012) and binds copper.

Further we asked what might be the regulator(s) that affects ATP7B’s recycling back from the lysosome. In a preliminary proteome analysis on GFP-ATP7B vesicles isolated from HepG2 cells (data not shown), we have identified members of the retromer complex. Previously (Harada et al., 2000) had shown that ATP7B resides in late endosome (Rab7 positive) in high copper. Retromer on the other hand is recruited on endosomal membrane by sequential action of Rab5 and Rab7 (Rojas et al. (2008). Further, Priya et al. (2015) dissected the interaction of Rab7 and retromer complex and demonstrated that Rab7 recruits retromer to late endosomes via direct interactions with N-terminal conserved regions in VPS35.

Before investigating the role of retromer in retrieval of ATP7B from lysosomes, we first determined if ATP7B is stable in lysosomal and Rab7 compartments. Interestingly it was reported by Polishchuk et al, that even up to 200uM copper treatment in HepG2 cells, ATP7B shows no significant degradation. However, we noticed a drop in ATP7B abundance indicating degradation at 250uM copper at 2 hrs though the cellular architecture apparently looks normal under phase contrast microscope. At 50μM copper, ATP7B did not exhibit any degradation.

Upon examining triple-colocalization of ATP7B, VPS35 and Lamp1 in fixed cells, we notice that the level of overlap is moderate at high copper. This might be attributed to the fact that at a given point in copper treated cells, the nature of vesicles are highly heterogeneous comprising of retrograde and anterograde vesicles and that too at various stages of trafficking. We hypothesize that the lysosomes would exhibit a higher co-residence of VPS35 and ATP7B if we are able to synchronize the TGN exit (upon copper treatment) and lysosomal exit (upon subsequent copper removal) of ATP7B. However, time lapse imaging showed that GFP-ATP7B and mCherry-Vps35 colocalizes in the compartment for a few minutes. In VPS35 kd cells, ATP7B is trapped in the lysosomes even upon activating the retrograde pathway (Copper>BCS). We reason that ATP7B recycles back to TGN from the lysosomes directly and not via plasma membrane as we did not observe ATP7B staining at the plasma membrane or even at the cortical actin (data not shown).

Interestingly, we did not detect any direct interaction of ATP7B and VPS35 (or VPS26). This is possibly due to the fact that though retromer complex regulates lysosomal exit of ATP7B, the interaction is mediated via a different member of the complex. This proposition is substantiated by co-detection of ATP7B, VPS35 and VPS26 in a supercomplex (>>250kDa) and also proximity of ATP7B and VPS35 at a range <40nm as detected in the PLA study. Retromer complex shows heterogeneity in its subunits that are responsible for binding to the cargo (Follett et al. (2016) Zhang et al. (2012) Belenkaya et al. (2008); Suzuki et al. (2019) Feinstein et al. (2011). It has been shown that the canonical recycling signal for the Divalent Cation Transporter (DMT1-II) binding of retromer is mediated via the interface of VPS26 and SNX3 in a hybrid structural model shows that the α-solenoid fold extends the full length of Vps35, and that Vps26 and Vps29 are bound to its two opposite ends (Lucas et al. (2016) Hierro et al. (2007). This extended structure suggests that multiple binding sites for the SNX complex and receptor cargo are present. It has been shown show that membrane recruitment of retromer is mediated by recognition of SNX3 and RAB7A, by the VPS35 subunit. These bivalent interactions prime retromer to capture integral membrane cargo, which enhances membrane association of retromer and initiates cargo sorting (Zhao et al., 2007). Further studies are needed to be carried out to identify the exact interface of ATP7B-retromer interaction.

How copper (or copper removal) mediates triggering of ATP7B’s retrograde pathway is not understood. It is likely that copper binding to the 6 MBD on ATP7B N-terminus, exposes the upstream 1-63 N-terminal domain containing the ^41^NXXY^44^ domain. However, role of C-terminus cannot be completely discounted as Braiterman et al has shown that multiple regulatory phosphorylation sites lie on the C-terminus that might play an indirect role in regulation of ATP7B by retromer complex (Braiterman et al., 2015).

Wilson disease, though a Mendelian disorder caused by mutations only in *ATP7B* gene, shows a large spectrum of symptoms and age of onset. We hypothesize that polymorphisms and mutations in trafficking regulatory proteins might be responsible for imparting such high phenotypic heterogeneity. Mutations and SNPs in the retromer subunit genes are associated with many hereditary conditions (Reitz (2018); Small (2008); Chen et al. (2017) Rahman and Morrison (2019); Shannon et al. (2014). Varadarajan et al, reported significant association of SNPs of retromer complex genes (SNX1, SNX3 and Rab7A) with Alzheimer’s disease Vardarajan et al. (2012) Similarly, VPS35 hemizygous condition accentuates Alzheimer’s disease neuropathology (Wen et al. (2011). Additionally, Parkinson’s disease-linked *D620N VPS35* knockin mice manifest tau neuropathology and dopaminergic neurodegeneration (Chen et al. (2019). It would be important to extend the knowledge of role of retromers in ATP7B trafficking to delineate genotype-phenotype correlationship in Wilson disease patients.

## Materials and methods

### Plasmids and reagents

GFP-ATP7B construct was available in lab. The mCherry WT-VPS35 and mCherry VPS35 (*R107A*) construct was kindly gifted by Dr. Sunando Datta, IISER Bhopal, India. pET28aSUMO and pGEX vectors was kindly gifted by Dr. Rahul Das, IISER-Kolkata, India.Following are the antibodies that has been used for experiments: rabbit anti-ATP7B (# ab124973), mouse anti-golgin97 (# A21270), goat anti-VPS35 (# NB 100-1397), mouse anti-VPS26 (# NBP 236754), mouse anti-VPS35 (# sc-374372); for western blot, mouse anti-Lamp1 (DSHB: # H4A3), mouse anti-Rab7 (# sc-376362), Donkey anti-Rabbit IgG (H+L) Alexa Fluor 488 (# A-21206), Goat anti-Rabbit IgG (H+L) Alexa Fluor Plus 647 (# A32733), Donkey anti-Mouse IgG (H+L) Alexa Fluor Plus 647 (# A32787), Donkey anti-Goat IgG (H+L) Alexa Fluor 568 (# A-11057), Donkey anti-Mouse IgG (H+L) Alexa Fluor 568 (# A10037). Endo toxin free plasmid isolation was done using EndoFree Plasmid Maxi Kit (# 12362).

### Cell lines and cell culture

HepG2 cells were grown and maintained in complete medium containing low glucose Minimum Essential Medium (MEM) (# 41500-034) supplemented with 10% Fetal Bovine Serum (# 10270-106), 1X Penicillin-Streptomycin (# A001), 1X Amphotericin B (# 15290026). Similarly HEK293T cells were grown and maintained in Dulbecco’s modified Eagle’s medium (DMEM) (# CC3004.05L) supplemented with 10% Fetal Bovine Serum,1X Penicillin-Streptomycin, 1X Amphotericin B. For transfection of plasmids in HepG2 cell, Lipofectamine 3000 reagent (# L3000-001) was used according to manufacturer’s protocol. For transfection in HEK293T, for live cell imaging, JetPrime (# 114-07) transfection reagent was used.

### Knockdown assays

Accell Human VPS35 (55737) siRNA-SMARTpool (#E-010894-00-0010), Accell Non-targeting siRNA (#D-001910-01-05), Accell siRNA Delivery Media (#B-005000-100), 5X siRNA Buffer (#B-002000-UB-100) and Molecular Grade RNase-free water (#B-003000-WB-100) were purchased from Dharmacon. HepG2 cells were seeded in complete medium at a density of 1.5×10^5^ cells/ml in coverslips heat fixed on 24-well plate. Cells were allowed to double for approx. 48hrs (doubling time of HepG2). After 48 hours, media was discarded, rinsed with 1X PBS pH 7.4 and si RNAs were added at a final concentration of 1μM resuspended in Accell siRNA Delivery Media (# B-00-5000-100). This condition was maintained for 72 hours after which the si-RNA containing media was replaced with complete media and again maintained for another 24 hours. This ensures better knock down at protein level. To validate knock down of VPS35, western blot was performed following same protocol from one well of 24 well plate.

### Immunofluorescence

HepG2 cells were seeded at a density of (0.8-1.6×10^5^ cells/ml) on coverslips heat fixed on wells of 24 well plate each time while conducting immunofluorescence. Any treatment was performed at a confluency of (60-70) %, including transfection. 4% Para-formaldehyde (PFA) fixation was done following treatment. After fixation cells were permeablized with chilled methanol and finally washed with 1X PBS. Fixation and permeablisation was carried out in cold condition. Cells were blocked in 3% BSA suspended in 1X PBS for either 2hours at room temperature (RT) or O/N at 4°C. Following this, primary antibody (1^0^) incubation was done at RT in moist chamber for 2hours. After 1^0^ incubation, cells were washed with 1X PBST for 3 times and again re-incubated with corresponding secondary antibodies (2^0^) for 1.3 hours at RT. This was followed by further washing with 1X PBST for 3 times and finally with 1X PBS for two times. Coverslips were fixed on glass slides using SIGMA Fluoroshield™ with DAPI mountant. (#F6057). The solvent for antibody suspension was 1% BSA in 1X PBST.

For STED sample preparation, HepG2 cells were seeded on glass coverslips, treated with BCS and copper (as mentioned in Fig. 8E). Treatment was done at 70% confluency. Cell were fixed with 2% PFA for 20mins followed by washing with 1X PBS, pH 7.2 for 15mins (× 2) and then quenched with 50mM NaCl. Blocking and permeation was done for 30mins with 1% BSA along with 0.075% saponin. Cell were co-incubated with primary rabbit anti-ATP7B and goat anti-VPS35 for 2hrs at room temp. Followed by 1X PBS washing and incubation with secondary anti-rabbit Alexa 488 and anti-goat Alexa 647. Coverslips was mounted with ProLong™ Diamond Antifade Mountant with DAPI (# P36962).

### Time-lapse fluorescence microscopy

HEK293T cells were seeded on confocal dishes (SPL) and were co-transfected separately with GFP-ATP7B and mCherry-VPS35-WT, mCherry-VPS35-MT(R107A) (gift by Sunando Datta, IISER Bhopal), using jetPRIME (Polyplus) transfecting regent as per manufacturer protocol. Images were acquired using Leica SP8 confocal setup with 63x oil objective. For ATP7B&VPS35-WT/MT, images were taken at every 1.964 s interval using Lightning by Leica.All the images were processed using Fiji and LASX software provided by Leica and videos were processed using Cyberlink Powerdirector.

### Co-purification assays and co-immunoprecipitation

#### Copurification

Composition of bacterial lysis buffer for N-term and C-term ATP7B: 50mM Tris-Cl, 50mM NaCl, 5mM EDTA, 10% glycerol (5% for C-term), ~1mM beta-marcaptoethanol, pH-8.0. Same buffer was used for the washing after incubation of lysates with beads. Composition of HepG2 cell lysis buffer:1X PBS buffer with 250mM sucrose, 1mM EDTA, 1mM EGTA, 1mM PMSF and 1X protease inhibitor cocktail. Same buffer was used for the washing after incubation of lysates with beads. Wt-C-term and Wt-N-term ATP7B were cloned into the pET28aSUMO and pGEX vectors respectively followed by transformation into competent BL21 E.coli for the bacterial expression of the proteins. BL21 containing Wt-N-term ATP7B was grown in Luria broth in presence of 100μg/ml ampicillin whereas BL21 containing Wt-C-term ATP7B was grown in nutrient broth in presence of 50μg/ml kanamycin followed by induction with 1mM isopropyl β-D-thiogalactopyranoside (IPTG) at 18°C and 37°C respectively for 16hr. BL21 and empty pGEX were used as negative control for C-term and N-term ATP7B co-purification respectively. Cells were resuspended in lysis buffer and lysed by sonication (100 amplitude/10 sec on/30 sec off) x 7 to 8 cycles. Bacterial lysates (collected from 100ml culture) were incubated with the Ni-sepharose (for C-term) and GSH Beads (for N-term) for 3hr at 4°C followed by washing using respective buffers. HepG2 pellet (collected from 10mm dish) was lysed by sonication (100 amplitude/10 sec on/30 sec off) x 5 cycles using respective lysis buffer for C-term and N-term co-purification. Insoluble materials were sedimented at 13,200rpm for 20mins at 4°C and supernatant was incubated with beads for 3hr at 4°C followed by elution with 3XSDS-PAGE loading buffer. Eluted products were used for western blotting using anti-VPS35 and anti-VPS26 antibodies.

#### Co-immunopreciptation

All solutions were pre-chilled to 4°C and all steps were carried out on ice. HEK293T cells were transfected with GFP-ATP7B and treated with different Cu conditions followed by washing with 1x PBS and lysis using lysis buffer (10mM Tris-Cl pH 7.5, 150mM NaCl, 0.5mM EDTA, 0.5 % NP40, PMSF and protease inhibitor cocktail in ddH_2_O). Cell extracts were triturated with 2ml syringe and incubated for total 45minutes and the insoluble materials were sedimented at 16,000*g* for 10 min at 4°C. Co-IP experiment was performed using GFP-trap beads (ChromoTek, # gta-20) following the manufacturer protocol. The supernatants were diluted using diluted buffer (10mM Tris-Cl pH 7.5, 150mM NaCl, 0.5mM EDTA, PMSF and protease inhibitor cocktail in ddH_2_O to yield 0.25 % NP40) and incubated with GFP-trap beads for 2hr at 4°C on a rotating wheel. Finally the interacting proteins were eluted using 0.2M glycine and used for western blotting. Western blotting of VPS35, VPS26 and GFP: Samples for Western blotting were resolved by sodium dodecyl sulphate-polyacrylamide gel electrophoresis (SDS-PAGE) and separated proteins were transferred to nirocellulose membrane. After protein transfer, the membrane was blocked in 5% non-fat milk powder (for VPS35&VPS26) and 3% BSA (for GFP) and incubated with primary antibody diluted in 5% non-fat milk powder (for mouse anti-VPS35 & mouse anti-VPS26 1:1500 dilution) or 1%BSA (for rabbit anti-GFP 1:10000 dilution) overnight at 4°C. Following incubation, the membrane was briefly washed three times with TBS-T and incubated with HRP-conjugated secondary antibodies (anti-mouse HRP 1:5000 dilution and anti-rabbit HRP 1:15000 dilution) diluted in 5% non-fat milk powder or 1%BSA for 1.5hr at RT. The membrane was washed three times for 5 min in TBS-T flowed by two times washing with TBS at RT and incubated for 5min at RT with Enhanced Chemiluminescence (ECL) substrate and ECL plus (1:1).

### Immunoblotting

HepG2 cells were grown on 60mm dish and cell pellet was collected at 70% confluency. For lysate preparation of membrane protein dry pellet was dissolved in 200μL of lysis buffer (composition: sucrose 250mM, EDTA 1mM, EGTA 1mM, 1X PBS as solvent, 1X protease cocktail inhibitor) and incubated on ice for 1hour with intermittent tapping. Dounce homogenization of dissolved pellet was done for approx. 400 times followed by syringe up down with 22-24 gauge needle for 20-25 times on ice. This enables the cell to rupture completely. The soup was centrifuged at 600 R.C.F at 4°C for 10mins to discard debris and nucleus. Further mitochondrial fraction was discarded by centrifugation at 3000 R.C.F for 10mins at 4°C. The resultant soup was subjected for ultra-centrifugation at 1,00,000 R.C.F for 1hour at 4°C to collect membrane fraction. Pellet was dissolved in membrane solubilizing buffer (composition: sucrose 250mM, EDTA 1mM, EGTA 1mM, NP-40 1.0%, Triton X-100 1.0%, 1X PBS as solvent, 1X protease cocktail inhibitor). For, soluble protein, whole cell lysate was prepared with RIPA lysis buffer (composition: 10mM Tris-Cl pH 8.0, 1mM EDTA, 0.5mM EGTA, 1.0% Triton X-100, 0.1% sodium deoxycholate, 0.1% sodium dodecyl sulphate, 140mM NaCl, 1X protease cocktail inhibitor). Dry pellet was dissolved in RIPA lysis buffer and incubated on ice for 30mins with intermittent tapping. The solution is then sonicated with a probe sonicator (3-4 pulses, 5sec, 100mA). Followed by this, centrifugation at 20,000 R.P.M for 20mins at 40C was done to pellet down cellular insoluble debris and soup was collected. Protein estimation was carried out with Bradford reagent (B6916-500ML) following manufacturer’s protocol. Protein sample preparation was done by adding 4X loading buffer (composition: Tris-Cl pH 6.81, 4% SDS, 10% β-ME, 20% glycerol, 0.02% bromophenol blue, urea 8M) to a final concentration of 1X and ran on SDS PAGE (6% for membrane fraction and 10-12% for soluble fraction) to separate proteins according to molecular mass. This was further followed by wet transfer of proteins onto nitrocellulose membrane (1620112, BioRad). After transfer, the membrane was blocked with 3% BSA in 1X Tris-buffered saline (TBS) buffer pH7.5 for 2hrs at RT with mild shaking. Primary antibody incubation was done overnight at 4° C following blocking and then washed with 1X TBST (0.01% Tween-20) for 10mins (× 3 times). HRP conjugated respective secondary incubation was done for 1.3 hrs at RT, further washed and signal was developed by ECL developer (170-5060, BioRad/ 1705062, BioRad) in chemiluminescence by Chemi Doc (BioRad)

### In-cell crosslinking and co-immunoprecpitation of crosslinked products

HEK293T cell was transfected with GFP-ATP7B and supplemented with modified DMEM-LM media (without leucine and methionine) and photo Leucine and photo Methionine. Cell was treated with either 100μM CuCl_2_ or 25μM TTM (tetrathiomolybdate) for 4hrs and subjected to UV crosslinking at 365nm for 15mins. Cell was processed for co-immunoprecipitation (Co-IP) assay following GFP-Trap_A (code-gta-20) Chrormotek with few modifications. All steps were carried out on ice. Cell lysis was done with Co-IP compatible 200 μl lysis buffer (10 mM Tris/Cl pH 7.5, 150 mM NaCl, 0.5 mM EDTA, 0.5 % Nonidet™ P40, 1X protease cocktail inhibitor and 1mM PMSF). Trituration was done with 26 gauge needle in 1ml syringe for approx. 40times for cell to completely lyse followed by centrifugation at 3000g for 5mins at 4°C. The sup was collected and diluted with 300 μl of dilution buffer (10 mM Tris/Cl pH 7.5, 150 mM NaCl, 0.5 mM EDTA). GFP-Trap A beads slurry was equilibrated with dilution buffer and incubated with the diluted supernatant and subjected to tumbling end-over on a rotatory wheel at 10rpm for 4hrs (first two hours at room temperature and the next two hours at 4°C).

### Proximity ligation assay

Proximity ligation assay was performed according to manufacturer’s protocol (*Duolink™ PLA Technology, Sigma)*. Briefly, HepG2 cell was grown on glass coverslips upto 70% confluency and treated with copper for 2hrs, fixed with 4% paraformaldehyde and permeabilized with 0.1% saponin. Thereafter, cell was blocked with blocking solution (DUO82007) for 1hr at 37°C and incubated with primary antibodies (as indicated in Fig.10) for 2hrs at 37°C followed by washing twice with Wash Buffer A (DUO82049-4L) for 5mins on gentle orbital shaker at room temperature (RT). During washing, the secondary PLA probes (DUO92005, DUO92003, DUO92001) were diluted to 1:5 with antibody diluent (DUO82008) and cell was incubated with it for 1hr at 37°C. Cell was further washed in same way. During washing, the ligation mixture was prepared by diluting the ligase enzyme (DUO82029) 1:40 in 1X ligation buffer (DUO82009) and applied onto cell. It was then incubated for 30mins at 37°C followed by washing twice in Wash buffer A for 2mins at RT. During washing, the amplification mixture was prepared by diluting the polymerase enzyme (DUO82030) 1:80 in 1X amplification buffer (DUO82010) and applied on cell with incubation at 37°C for 100mins. After incubation, cell was washed twice with Wash Buffer B for 10mins. A final wash with 0.01X Wash buffer B was given prior mounting with DAPI containing mountant. Images was acquired with Leica SP8 confocal platform using 576 nm (Cy3) filter.

### Microscopy

All images were acquired with Leica SP8 confocal platform using oil immersion 63X objective and deconvoluted using Leica Lightning software. For Structured Illumination Microscopy, images acquisition was taken at 100X magnification in Zeiss Elyra PSI. For Stimulated emission depletion (STED) microscopy, imaging was done in Leica STED 3X. For Alexa 647, 775 STED laser line was used and for Alexa 488, 592 laser line was used for depletion. Line average was set at 4 and pixel size was kept as 25nm to achieve maximum resolution. STED corrected images were deconvoluted and processed by Scientific Volume Imaging of Huygens Professional Software with default settings.

### Image analysis and statistics

Images were analyzed in batches using ImageJ (Schneider et al. (2012), image analysis software. For colocalization study, Colocalization_Finder plugin was used. RO Is were drawn manually on best z-stack for each cell. For three protein colocalization study, the other two protein co-residing vesicles were isolated using Analyze Particle tool, and colocalization study were carried on with the reference protein, ATP7B in our case. RGB_Profiler plugin was used to obtain the line profile graph. For statistical analysis and plotting, ggplot2 (Wickham (2009) package was used in R v-3.4.0 (Team (2015). Non-parametric tests for unpaired datasets (Kruskal Wallis test and Mann-Whitney U test) were performed for all the samples.

## Supporting information

1B: ATP7B and R107A-VPS35

1A: ATP7B and wt-VPS35

## Online Supplementary materials

Video: Time lapse imaging to record colocalization of ATP7B (green) and VPS35 (red) in high copper. 1A: ATP7B and wt-VPS35; 1B: ATP7B and R107A-VPS35

## Acknowledgments

This work was supported by Wellcome Trust India Alliance Fellowship (IA/I/16/1/502369) and Early Career Research Award (ECR/2015/000220) from SERB, Department of Science and Technology (DST), Government of India and IISER K intramural funding to AG. SM and IB was supported by Pre-doctoral fellowship from Council of Scientific and Industrial Research, India. TS was supported by National Postdoctoral Fellowship, SERB, India. We thank Dr. Ashima Bhattacharjee for critical review of the manuscript.

## Author contributions

AG and SD designed the experiments and wrote the manuscript. SD, Ruturaj and TS did the experiments and analyzed the data. SM wrote the codes and analyzed the data. Ruturaj helped with time-lapse imaging. IB conducted the experiments with the ATP7B mutants. Dr. Anupam Banerjee (Zeiss) helped us with SIM imaging at JNCASR. STED microscopy was carried out at the Leica microscopy facility at IISER Pune. All authors reviewed the results and approved the final version of the manuscript.

We thank Rahul Das (IISERK), Arindam Mukherjee (IISERK), Oishee Chakrabarti (Saha Inst of Nuclear Physics), Ashima Bhattacharjee (Amity Univ. Kolkata) for sharing their reagents and instruments with us.

The authors declare no competing financial interests.

## Legends to supplementary figures

**Fig S1:**
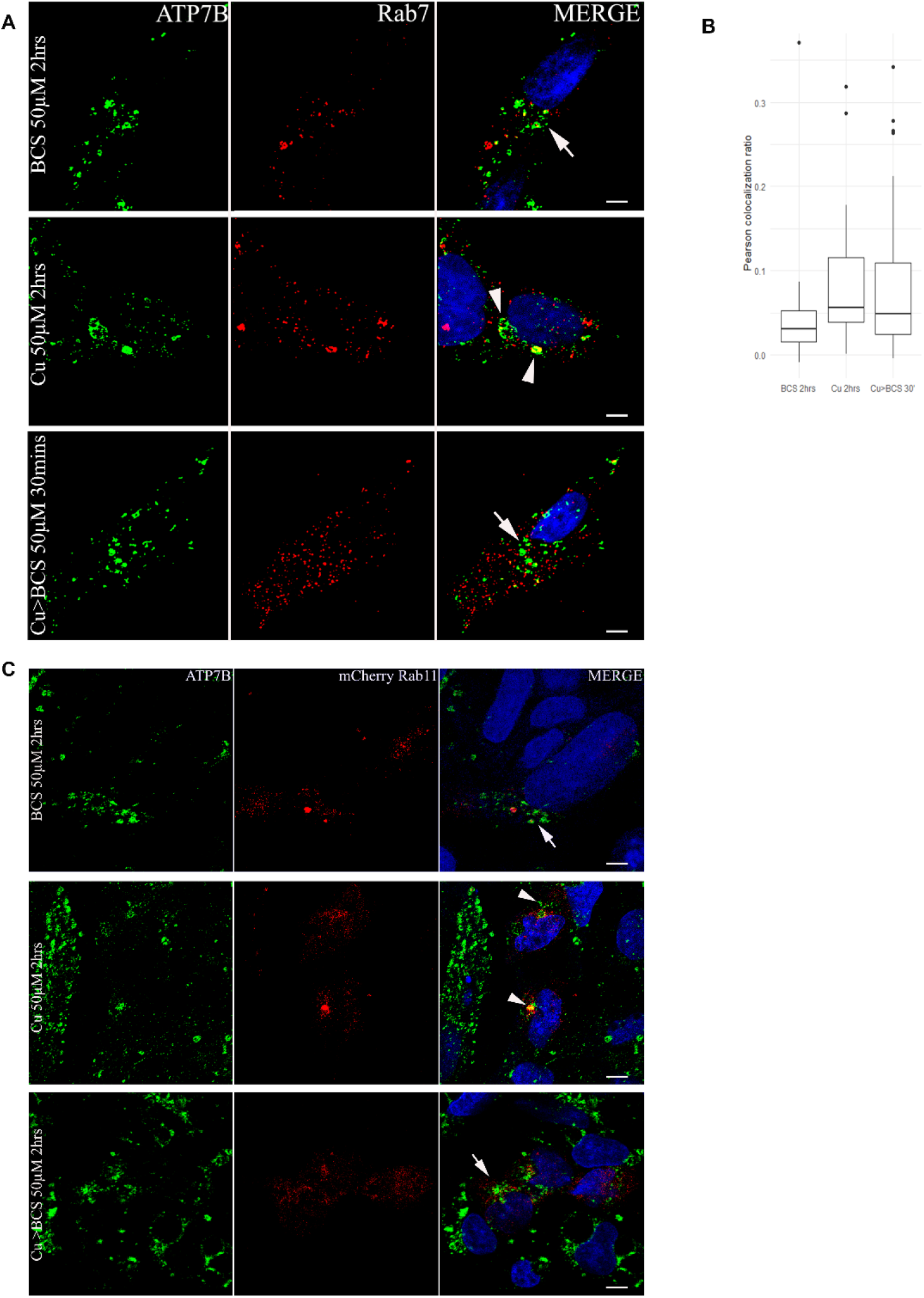
Colocalization of ATP7B with late (Rab7) and mid (Rab11) endosomal markers at different copper levels: (A) Colocalization of ATP7B (green) with late endosome marker, Rab7 (red) in copper limiting, BCS (top panel) and 50uM copper (panel 2) and copper depletion post copper treatment (bottom panel). Scale bars represents 5μM. Blue signal represents DAPI staining for nucleus. (B) Pearson’s correlation coefficient of colocalization between ATP7B and Rab7 at different copper conditions demonstrated by a box plot. (C) Colocalization of ATP7B (green) with mid endosome marker, Rab11 (red) in copper limiting, BCS (top panel) and 50uM copper (panel 2) and copper depletion post copper treatment (bottom panel).

**Fig S2 (A):**
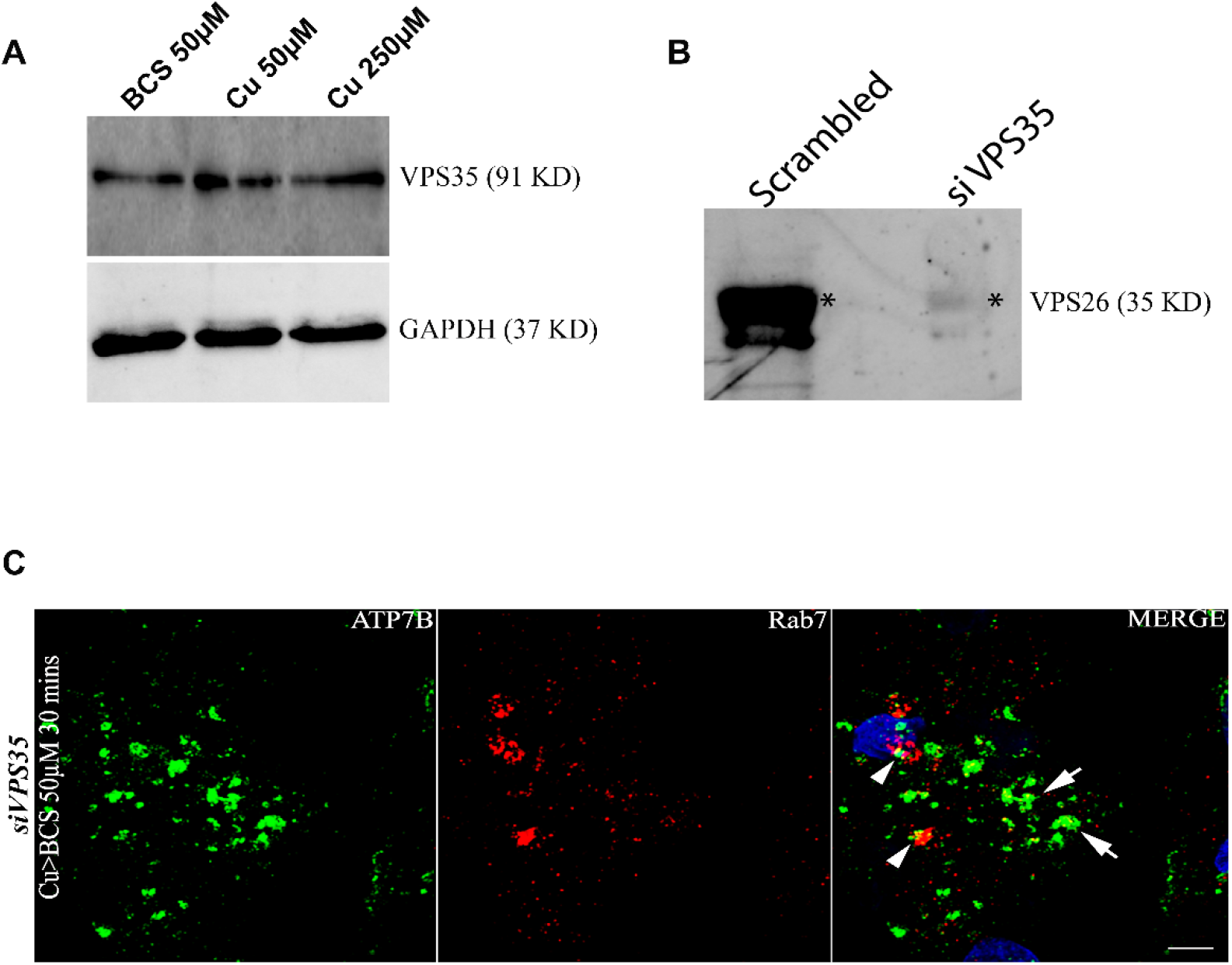
Immunoblot of VPS35 in different Copper conditions: Upper panel shows abundance of VPS35 remain unchanged in all copper conditions. Lower panel shows the same for GAPDH, as a control for cytosolic proteins. (B) **Immunoblot of VPS26 in siVPS35 HepG2:** siRNA mediated knockdown of Vps35 in HepG2 cells shows downregulation of its core partner VPS26 as compared to its control. (*) denotes the VPS26 protein. (C) **VPS35 regulates retrieval of ATP7B from late endosome to TGN:** siVPS35 treated HepG2 cell shows colocalization of ATP7B (green) with Rab7 (red) in copper depletion post copper treatment for 30mins. Arrowhead shows ATP7B located in late endosomal vesicles and arrows shows non-returning vesicularized ATP7B. Scale bars represent 5μM. Blue signal represents DAPI staining for nucleus.

**Fig S3:**
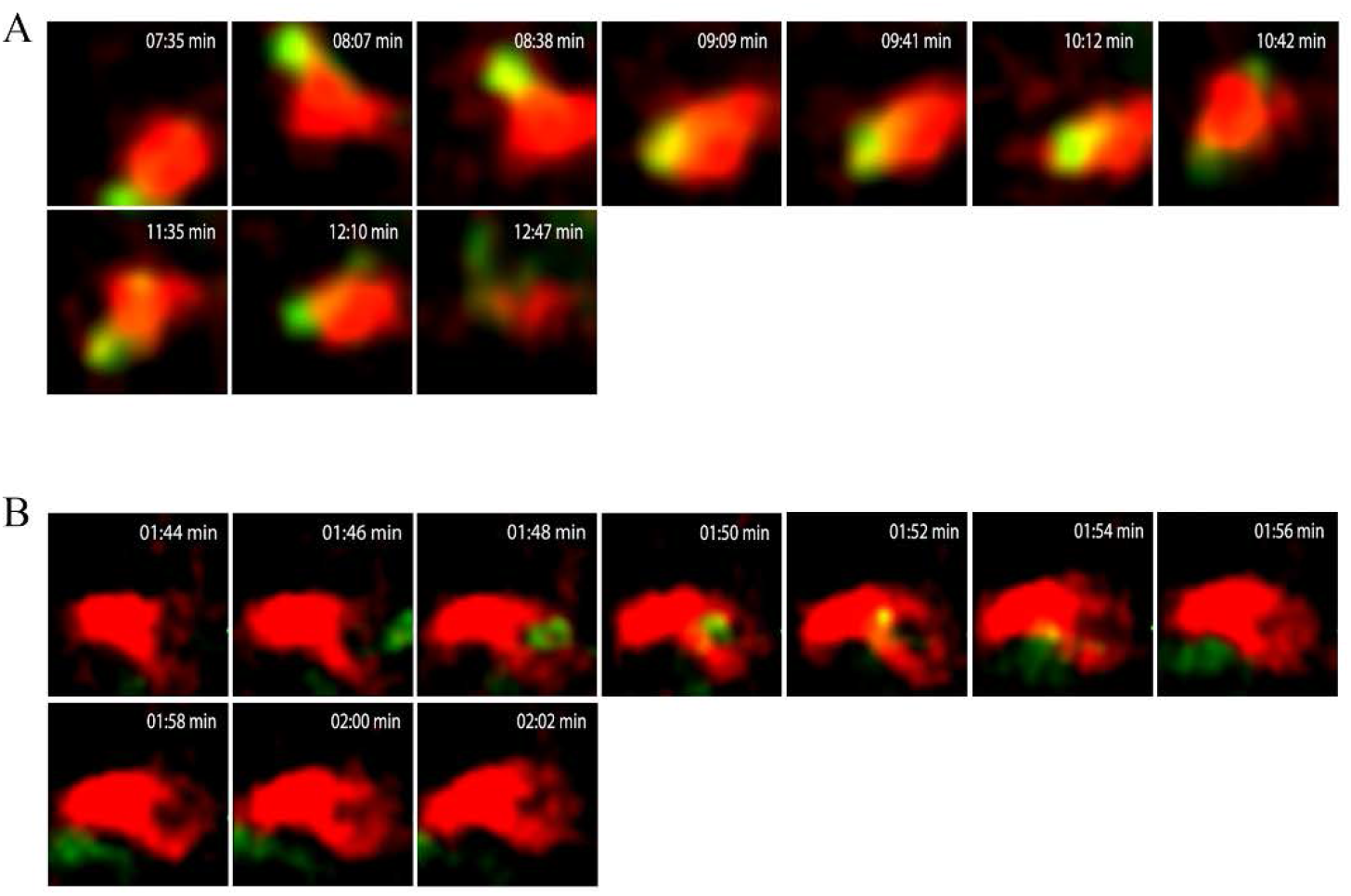
Comparative dwell time analysis of ATP7B and wtVPS35 vs its mutant, VPS35 R107A: Live-cell time-lapse high resolution deconvolution confocal microscopy shows dwell time of GFP-ATP7B with mCherry VPS35-WT to be ≈ 4 mins (A) and for mutant ≈ 3 seconds (B) in 50uM copper. Images were taken at every 1.93 s interval.

**Fig S4 (A):**
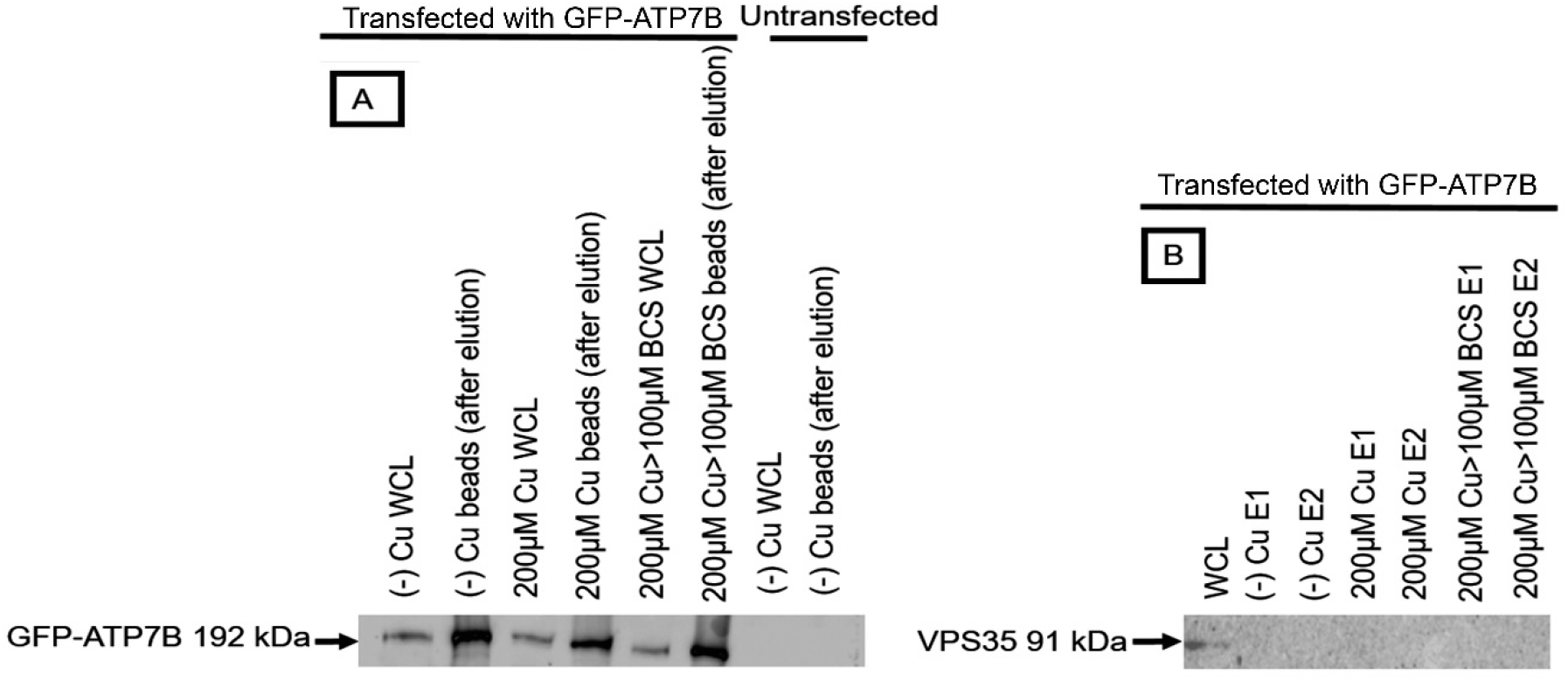
(A) Co-immunoprecipitation assay to determine the interaction between full length ATP7B and VPS35: Cell was transfected with GFP-ATP7B and treated with different copper conditions as mentioned. Lysates were incubated with GFP-trap beads. Untransfected cells were used as negative control. A) Immunoblot showing the presence of GFP-ATP7B in whole cell lysates and GFP-trap beads after elution. B) GFP-trap immune-co-precipitated products were subjected to immunoblot as indicated. Abbreviations: *FT: Flow through; WCL: Whole cell lysate, E: Eluate*.

